# Self-Authenticating Genomic Materials with Advanced Genome Signatures

**DOI:** 10.1101/2024.11.02.621633

**Authors:** Zhaoguan Wang, Jingsong Cui, Qian Liu, Jiawei Li, Chi Guo, Yiyang Liu, Changyue Jiang, Hui Xue, Jiaojiao Li, Yonggang Ke, Hao Qi

## Abstract

The authenticity and integrity of synthetic genomic materials containing valuable intellectual property are essential for advancing scientific knowledge and enhancing biosafety. Existing DNA tags and watermarks, however, have limited efficacy due to low mutation tolerance and inadequate digital encoding capacity. Here, we present “Genome Signature”, a biochemically stable and tamper-resistant DNA labeling system that enables the creation of self-authenticating genomes. Central to this system is a novel Golomb-ruler-derived Genome-Comb, which efficiently maps extensive nucleotide sequences onto limited codons within endogenous genes, significantly improving error correction and data encoding across millions of nucleotides. Our method successfully recorded a 4.5 million-nucleotide genome in living *E. coli*. The Genome Signature autonomously identifies and corrects mutations and effectively encodes data within codon orders, ensuring genome integrity and authenticity. Furthermore, it allows precise tracking of sequences across different cells, potentially revolutionizing the development of reliable genomic materials in synthetic biology.

## Introduction

Synthetic genomic materials are gaining increasing significance in scientific research and industrial applications, particularly in fields such as biomedicine(*1*), biomanufacturing(*2*), environmental bioremediation(*3*) and agriculture(*4*) (**Fig. S1a-c**). The rapid advancement of genome synthesis and editing technologies has enabled the global production of synthetic sequences with diverse gain-of-function designs at an unprecedented pace. Synthetic genetic materials are now ubiquitous in many biomedical products, such as vaccines(*5*) and cell therapies(*6*). However, alongside this excitement, societal concerns have also emerged. Notably, the unauthorized use of cells and their genomic materials, often with high commercial value, remains a significant issue that continues challenging the biomedical and biomanufacturing industries(*7*). Additionally, the widespread use of GMOs with synthetic sequences places significant pressure on natural biological systems through horizontal gene transfer(*8, 9*), leading to genomic contamination. In these misuse, abuse, and biocontamination scenarios, genomic materials lack no definitive references for accurately determining their identity and authenticity. As products with both high industrial value and biological risks, synthetic genomes should be placed under close administration(*10*), which demands rapid and accurate determination of the authenticity and provenance of genomic materials found in products.

Synthetic DNA tags (e.g., DNA watermarks) are emerging as effective tools for labeling sequences of interest, thereby facilitating the identification, specification, and protection of intellectual property rights. For instance, watermark DNA sequences have been incorporated into synthetic genomes of organisms such as *M. mycoides*(*11*), *E. coli*(*12*), and yeasts(*13*). However, conventional DNA tags face several significant limitations. While they are useful for genome identification, they do not independently verify functionality or detect subtle changes within genomes. Their typically limited size restricts their capacity to carry information, and the introduction of extensive tag sequences into synthetic genomes can result in undesirable alterations to genome size and function. Furthermore, because synthetic DNA tags lack biological function, they are highly vulnerable to mutations during cell replication. Continuous and unpredictable mutations can lead to the loss of information encoded in DNA tags, even with existing error-correction methods (**Table S1**)(*14-16*). As cells proliferate, the accumulation of mutations across the genome(*17*) further undermines the accuracy and effectiveness of DNA tags in verifying genome authenticity and integrity.

Given the critical need to safeguard and track synthetic genomic materials(*10*), establishing new paradigms for the protection of synthetic genomes is essential. To meet this challenge, we propose developing an internal genomic signature (**Fig. 1a**) analogous to the digital signatures used in modern electronic documents and transactions. Digital signatures allow for independent verification of electronic documents and transactions without the need for centralized authority, a feature that is vital for optimizing the efficiency of large-scale operations. Similar principles of efficiency and security have driven the success of several groundbreaking technologies, such as digital signatures, the Diffie-Hellman key exchange (DHKE) algorithm, and Quick Response (QR) codes. These technologies have achieved widespread adoption by leveraging their core strengths in efficiency and security, enabling them to scale to monumental levels.

**Figure 1.**
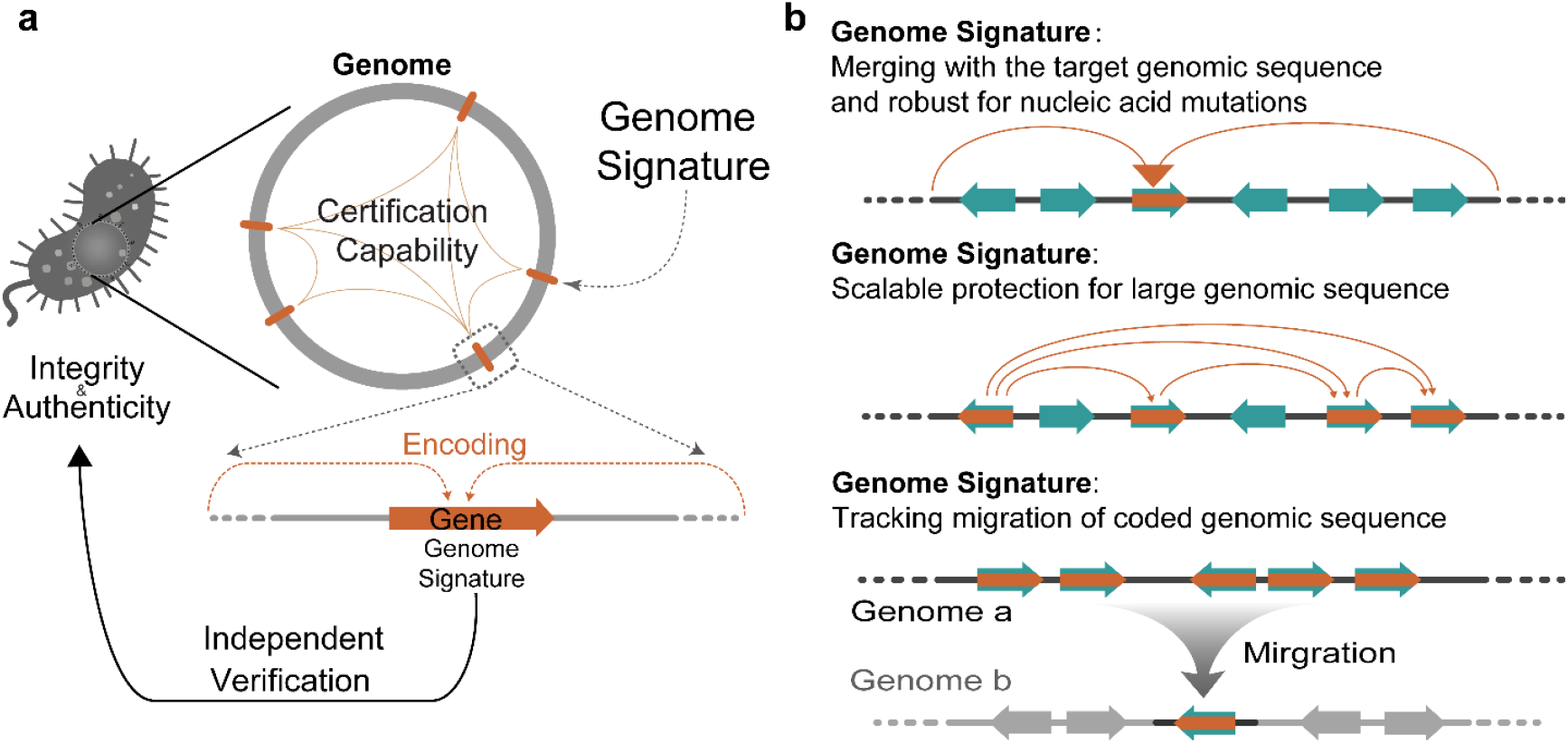
Genome Signature. Diagram illustrating the implementation and functionality of the Genome Signature in synthetic genomic materials. **(a**) Overview of the Genome Signature’s encoding process within a gene. The Genome Signature integrates into the target genome, providing certification capabilities that ensure integrity and authenticity. This enables independent verification of genomic sequences. (**b**) Functionalities provided by the Genome Signature. Integration with the target genomic sequence enhances robustness against nucleic acid mutations; Scalable protection mechanisms applicable to extensive genomic sequences; Tracking and verification of genomic sequence migration from Genome a to Genome b, ensuring accurate monitoring and protection of coded genomic sequences.

In this study, we developed a Genome Signature that allows for quick and accurate verification of the integrity and authenticity of individual genomic materials without the need for comparison to reference sequences (**Fig. 1b**). The Genome Signature is seamlessly integrated into the genomic sequence using Genome-Comb Hash Mapping (Genome-CHM), a specialized algorithm based on a Golomb-ruler-derived Genome-Comb. Unlike conventional DNA tags that merely label genomes, this method encodes nearly unlimited nucleotide sequences into the limited codon order of endogenous genes, preserving the essential and inseparable nature of the genome without altering its overall size. The Golomb ruler’s unique variable-length encoding scheme enables efficient computation across millions to billions of nucleotides. The Genome Signature enables the identification and correction of hundreds of random nucleotide mutations across *E. coli* genome, allowing for the verification of its integrity and authenticity without the need for reference sequence comparisons. Additionally, the signature-secured sequences remain identifiable even when inserted into different genomes. The Genome-CHM algorithm, with its great scalability and flexibility, provides robust self-quantification capabilities, offering comprehensive authentication-key protection, tamper-proof security, and rigorous certification protocols. Surpassing current DNA tags, this approach enables the direct documentation and identification of the uniqueness of synthetic genomic materials in a blockchain-like manner.

## Results

### Genome-CHM algorithm for Genome Signature

For devising a Genomic Signature with advanced certification capabilities, we face the challenge of creating a new algorithm that can efficiently handle large amounts of information and provide error correction at the scale of the entire genome sequence. First, instead of nucleotide, we encoded the codons of the open reading frames (ORFs) of intrinsic genes in a genome, so that the biological function of the gene can effectively protect it from being lost during cell replication. The genetic code is redundant and 18 of the 20 proteinogenic amino acids are encoded by multiple synonymous codons. Synonymous codon substitution is a major genetic polymorphism in nature that influences bacterial gene expression and is generally not fatal to the cell(*18*). Therefore, in addition to the nucleotide sequence, codon order is another information dimension hidden in the genome. However, to use codon order as an information-carrying medium, one hurdle is the non-uniform storage space due to the uneven number and frequency of synonymous codons for each amino acid. Another issue is that the number of codons in the genome is several orders of magnitude smaller than the number of nucleotides, making codon order comparably limited in terms of storage space. Therefore, even though codon redundancy was used to store data in synthetic genomes, only very simple message data, 498 bits encoding 83 words in gene T7 RNAP as the largest reported case(*19*), were encoded in previous studies (**Table S2**). However, beyond data encoding, we face an even greater challenge: embedding error correction codes into codons to ensure strong tolerance to mutations across the entire genome sequence, which comprises millions of nucleotides.

We developed a highly integrated algorithm Genome-CHM, incorporating modified Mixed Radix Coding (**mMRC**) (**Note S1**) and Directed Hash-Adaptive Codon Mapping (ACM) (**Note S1**), to encode both authenticity data and mutation correction codes for genome integrity information into gene codons (**Fig. 2a**). First, the authenticity data, including identity information of the genomic materials as well as the identity and length of the target gene, are encoded into the codons at the 5’ terminus of the gene, referred to as the Data Tag (DT), using a modified algorithm based on the Mixed Radix Coding (MRC) system we previously developed(*20*) (**Note S2**). Generally, each codon is assigned with one integer range of [0,60) according to the codon mapping table (**Table S3**) and the total number of all redundant codons for the same amino acid is fixed at 60. For instance, [0,20) is assigned to GAG (0.33 codon frequency) and [20,60) to GAA (0.68 codon frequency) in *E. coli*. Therefore, codon frequency in encoded genes can be controlled by adjusting the integer range assigned to each codon. In this study, unless otherwise stated, the frequency in coded genes is close to the natural frequency according to the codon mapping table (**Fig. S3b**).

**Figure 2.**
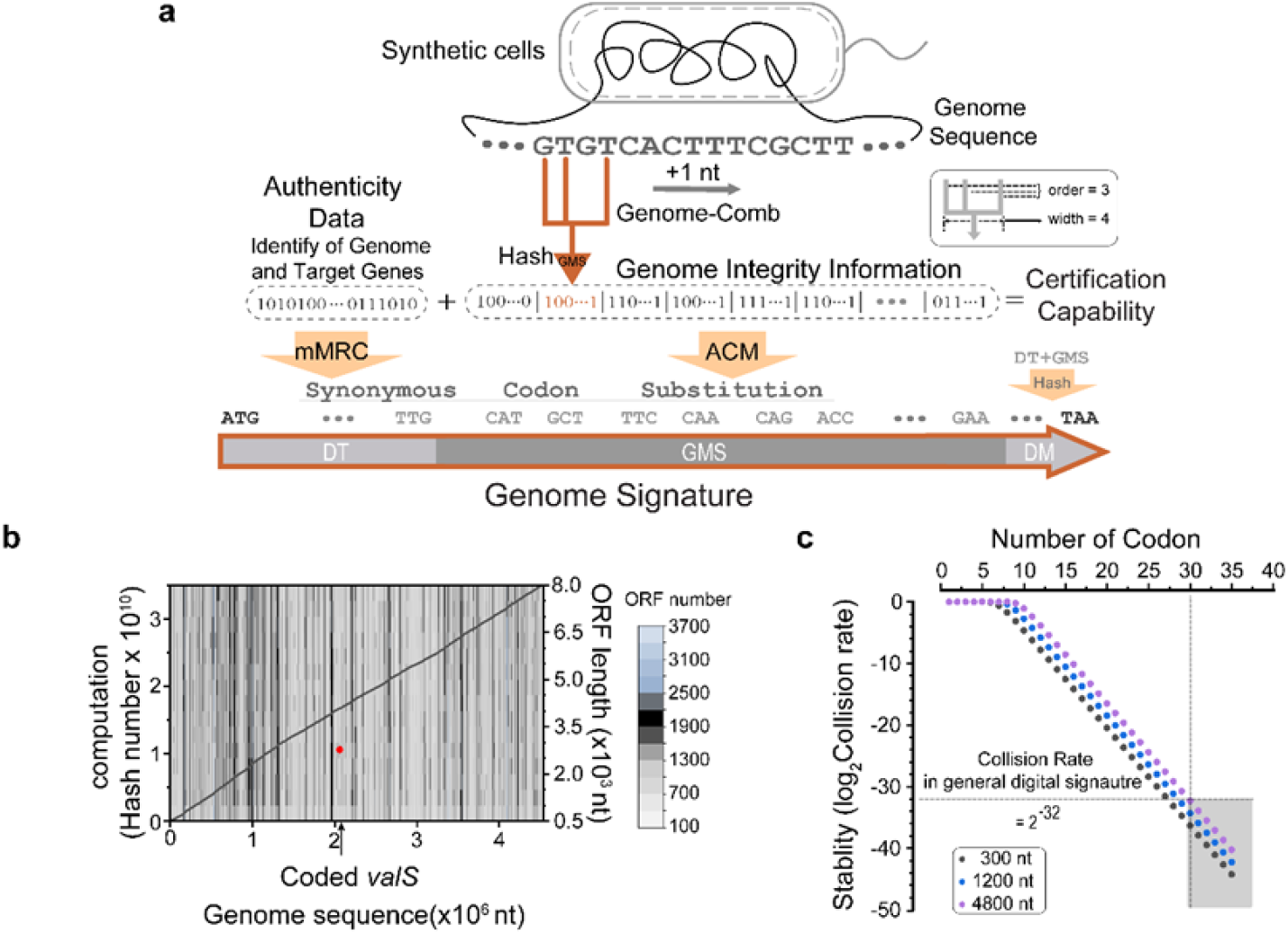
Genome-CHM algorithm. (a) Schematic representation of the Genome-CHM algorithm for encoding Genome Signature into synthetic genome sequence. The Genome-Comb mechanism generates a unique hash from scanning the genomic sequence, integrating it into the genome to include information on authenticity, identity, and integrity (**Note S1-S4, Supporting Information**). This enables certification capabilities by mapping genome integrity information and identifying any genetic modifications. (b) Computational assessment showcasing the relationship between yielded hash numbers and coded genome sequences. Identification of the coded *valS* takes place over 16 million ORF-like sequences (gray dot), each consisting of over 180 codons, found within the 4,521,562 nt *E. coli* genome sequences. This process involves approximately 3.5E+10 hash calculations (black line), and the coded *valS* location on the genome indicated by an arrowhead. (c) The stability of the Genome Signature is conducted by evaluating the collision rate of DM decoding computations in relation to the number of codons in the DM of an encoded gene. The collision rate of 2^-32^ (dashed line), which is the accepted level for digital signatures in electronic documents, is achieved for DM with 30 codons.

The relevant data are encoded into corresponding integers, which are then mapped to synonymous codons using the mMRC algorithm, rather than simply being converted into a nucleotide sequence. During encoding, the binary data is segmented, and each segment is mapped to a codon at a specific position within the target nucleotide sequence. The selection of synonymous codons at each position is constrained by factors such as optimizing codon usage and avoiding the formation of undesirable DNA fragments. At each step, a context-specific mapping table is generated to dynamically determine the appropriate codons, excluding those that may violate biochemical or preset constraints. This position-by-position encoding process continues until the entire binary sequence is converted into a synonymous codon sequence. The flexibility in adjusting the synonymous codon (radix) at each position enables high-density encoding while ensuring sequence stability and compatibility with degenerate or composite base schemes. In the current setting, the number of codons within a single gene, as well as the number of genes that can be encoded either together or separately, is effectively limitless. This allows for the flexible expansion of the information-carrying capacity (**Table S4**). Running on our desktop computer with a medium configuration (**Methods**), coding DT in essential gene *valS* with 952 codons was completed within seconds.

Next, we developed a novel algorithm to embed mutation correction codes for protecting genome integrity and to map these codes onto gene codons. This enables independent identification of mutations across the entire genome sequence and verification of integrity without the need for comparison to a reference sequence. However, the algorithm must encrypt large sequences, reaching millions of nucleotides, and its complexity significantly exceeds the capabilities of current methods. We achieve our ambitious goal by reimagining the Golomb ruler, an optimal mathematical structure that has previously been utilized in the fields of radio astronomy and nuclear magnetic resonance spectroscopy(*21*). This innovative variable-length encoding scheme has enabled us to create a special Genome-Comb (**Fig. 2a**). The Genome-Comb is capable of generating quantitative information for sequences of nearly unlimited length. This information records the relative position and content of each nucleotide, which is then mapped to codon order in a specific region within the gene. This mapping is achieved through a combined directed Hash function, commonly referred to as the GMS (Genome Mapping Storage). The GMS encompasses hundreds of codons located behind DT in the middle of the gene. Genome-Comb is designed to efficiently identify mutations across extensive sequences, including those that may span millions of nucleotides.

The Genome-Comb scans the genome sequence step-by-step (**Fig. S2**), with the adjustable parameters of ‘order’ representing the number of teeth and ‘width’ defining the largest distance between them. Within each step, a single strand of nucleotides, equivalent to the order of the DNA comb, is scanned into the Hash function *Hash*_*GHC*_ (*N*_*c,i*_ to produce a monomial, *GP*_*i*_ (*x*). In this monomial, the degree number represents one codon position in the GMS, and the coefficient corresponds to the value stored at that position (**Note S3**). The mathematical link between a nucleotide strand from Genome-Comb and a codon in the GMS is defined as the Comb-Hash mapping. Finally, all the monomials are added up via the polynomial operations *GP*_*∑*_(*x*), and the total summed coefficient is converted to one [0,60) number by modulo 60 and then mapped to one synonymous codon by ACM to generate the GMS sequence.

In the final coding stage, the last 30 codons before the stop codon are encoded as DM (Decoding Marker) using ACM (**Fig. S3a**). Generally, the nucleotides sequence in DT, GMS, and the last 30 amino acid sequences are firstly encoded to a polynomial *DP*_*i*_(*x*) one by one by the function of Hash_DM_ (**Note S4**). Next, all of them are summed up via polynomial operations of *DP*_*∑*_(*x*) and converted to one [0,60) integer by modulo 60 and then mapped to synonymous codons by ACM to generate the final codon sequence of DM. Thus, DM can not only verify the authenticity and integrity of the coded gene sequence but also effectively correct nucleotide mutations within the sequence. Unlike a simple genome tag, the Genome Signature of the coded gene is algorithmically merged with the entire genome sequence. Its essential biological function leads to robust stability, making it resistant to being easily lost or rapidly changed during evolution.

Distinct from current frameworks, Genome Signature offers a certification capability akin to a digital signature, ensuring independent verifiability and enhanced security. Firstly, the coded gene can be independently identified from the raw sequencing data of the genome, eliminating the need for reference comparison. Generally, the verification of whether an ORF sequence is a coded gene is done by examining its DM (**Fig. S4a, b**, and **Note S4**). The ORF sequence is subjected to generate one new 30-codon string as DM′. If DM′ matched with the current last 30-codon sequence, then this ORF is one coded gene. Furthermore, the DM can tolerate the mutation within the ORF sequence. If the DM′ is not matched, each nucleotide in the ORF sequence is revised to all potential substitutions, deletions, or insertions one by one. Each revised sequence is then subject to DM decoding. The cost of DM mutation correction, measured in the number of Hash calculations, increases with the number of codons. Theoretically, the DM algorithm can correct multiple random mutations, but at the cost of increased computing resources (**Note S5**).

### Validation of Genome Signature encoded in the *E. coli* genome

The unique mathematical properties of the Golomb ruler-derived Genome-Comb enable the independent verification of the integrity and authenticity of the entire genome sequence. Without requiring sequence comparison, mutations—along with their precise locations and original nucleotide contexts—can be directly identified within the nucleotide sequence from raw genome sequencing data. The procedure for the independent identification of Genome Signature from the entire genome sequencing data is illustrated in **Figure S4**. Each ORF-like sequence in the genome, consisting of over 180 codons from ATG to any stop codon (TAA, TAG, or TGA), is individually subjected to DM decoding. As a result, over 16 million Genome Signature candidate sequences have been decoded in this *E. coli* genome (**Fig. 2b**). It took a few minutes to identify the Genome Signature *valS* after approximately 7E+10 hash calculations of all the candidate sequences (**Note S6**), running on our desktop computer with a medium configuration (**Table S5**). Although, the computation cost of mutation correction in DM decoding increases with sequence length, an O(n^e^) time-complexity (**Note S5**), the function of essential genes confers high sequence stability, generally making DM decoding with correction of single nucleotide mutations sufficient for identifying coded genes from sequences comprising millions of nucleotides or more.

The security of Genome Signature was accessed by quantifying the collision rate in DM decoding (**Fig. 2c**). For Genome Signature, a collision can be considered as a sequence with random mutations that generates the same hash value when encoding DM compared to the original sequence (**Note S7**). The collision rates are relatively low, and they decline sharply with an increase in the number of codons in DM. The collision rate for DM with 30 codons can be as low as 2^-32^, which is the widely accepted standard level for digital signatures in electronic documents.

For mutation identification at Genome, the genome sequence is encoded to generate one new codon order as GMS′ for comparison with the current GMS in the codon sequence of an identified coded gene (**Fig. 3a**). Any nucleotide mutation leads to multiple codon-changes in GMS′. Each codon-change in GMS′ can be traced back to multiple nucleotides using the corresponding Comb-Hash mappings. Each nucleotide can then be assigned a QA-score (questionable authenticity score) by counting all Comb-Hash mappings that point to it (**Note S8**). Theoretically, based on the special mathematical feature of the Golomb ruler(*21*), a mutated nucleotide in a sequence will be scanned “order” times by Genome-Comb and causes no more than 2×order codon-changes in GMS′. In GMS′, one nucleotide mutation leads to the new Comb-Hash mappings (error-in) and the loss of existing Comb-Hash mappings (error-out) (**Fig. S5a**). The error-in Comb-Hash mappings exclusively point to the mutated nucleotide, resulting in a higher QA-score. Conversely, the error-out Comb-Hash mappings lead to an increased number of codon-changes within GMS. However, the strong hash collision results in all codon-changes related to Comb-Hash mappings, except for error-in mappings, pointing towards nucleotides dispersed throughout the entire genome sequence. This leads to a relatively low QA-score noise. As shown in **Figure 3b**, all the correct nucleotides in the genome with a single coded gene, *valS* (an essential gene composed of 952 codons located at 3,900,299 - 3,903,154), have a score of zero. Conversely, a nucleotide with a high QA-score either indicates a mutation or is located near a mutation. As shown in **Figure 3c**, a nucleotide substitution (3,302,560 C>G) obtained a significantly higher QA-score compared to all other 4,521,561 nucleotides. Approximately 64.19% of all nucleotides scored zero, while over 99.99% of nucleotides had QA-scores lower than 0.3 (**Fig. 3d**).

**Figure 3.**
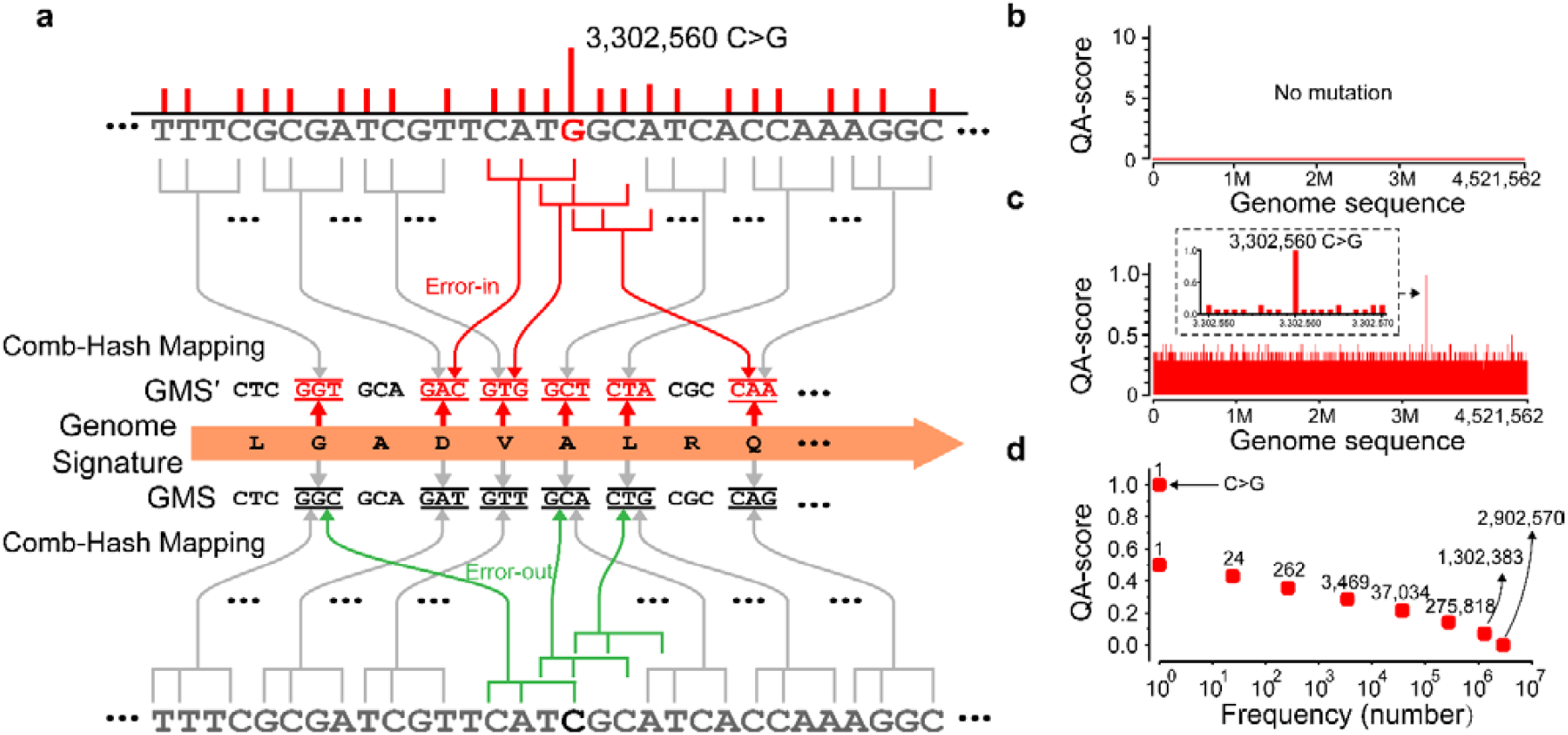
QA-score for mutation correction. (a) Diagram of the Comb-Hash Mapping process in GMS decoding illustrating the detection of a specific mutation (3,302,560 C>G). Substitution of 3,302,560 C>G causes changes in GMS′. Comb-Hash mappings of both the origin nucleotide C related (error-out, green line) and the substitution G related (error-in, red line) are quantified as a value, termed as QA-score (questionable authenticity score). Non-changed Comb-Hash mappings are indicated with gray solid lines. (b) Quality Assurance (QA) score profile across the genomic sequence showing baseline values in the absence of mutations. (c) QA score profile revealing a significant spike at the mutation position of substitution 3,302,560 (C>G) within the genome sequence, the inset displays the 20 nucleotides surrounding mutation. (d) The mutated C>G 3,302,560 nucleotide scored 1. All other nucleotides scored ≤0.5, and over 99.99% of them scored below 0.3.

Once the coded gene has been identified, the integrity of the entire genome sequence is evaluated using the QA score. This assessment facilitates the independent correction of nucleotide mutations throughout the genome. Specifically, all nucleotides in the genome are ranked by QA score in descending order and then subjected to revision—by substitution with one of the other three bases, deletion, or insertion of A, T, C, or G—until the GMS’ aligns with the GMS (**Fig. S5b)**. To demonstrate the accuracy of the QA-score, every nucleotide in the *E. coli* genome was mutated one-by-one. Over 99.9% of the mutated nucleotides were correctly identified by the GMS with high QA-scores, ranked in the top 1000 of QA-scores (substitution mutation nucleotides: 4,514,962, insertion mutation nucleotides: 4,515,469, and deletion mutation nucleotides: 4,514,867). The remaining mutated nucleotides were still ranked within the top 1% of high QA-score (**Fig. S5c**). In the GMS algorithm, the sensitivity to substitution is related to the number of Genome-Comb teeth, and the width for indels (**Note S3**). In the GMS algorithm, the sensitivity to substitution is determined by the number of Genome-Comb teeth, which refers to the pattern-matching capability of the algorithm. A higher number of Genome-Comb teeth leads to increased sensitivity and accuracy in detecting substitutions. Additionally, the width of Genome-Comb influences the sensitivity to indel mutations. A broader width for indels allows the algorithm to detect larger insertions or deletions more effectively. Therefore, GMS is efficient for identifying all types of nucleotide mutations (**Fig. S6**), and it further demonstrates that the Genome-Comb based QA-score is highly efficient in locating and correcting mutations in the whole genome comprising of over millions of nucleotides. The decoding cost of the GMS algorithm can be simply considered as a function of the total sequence length. This means that as the length of the input sequence increases, the decoding cost will also increase in a linear manner. This linear time complexity is referred to as O(n), where n represents the length of the input sequence. This aspect of the GMS algorithm makes it computationally efficient and scalable for large-scale mutation detection tasks (**Note S9**). For one substitution mutation correction, a total of 142,524 hash calculations were performed. These calculations were necessary to correct a single substitution in a genome containing a single coded gene of *valS*. In long-term bacterial evolution(*22*), it has been observed that single nucleotide mutations account for over 85% of genetic changes. Specifically, substitution mutations were found to occur almost 8 times more frequently than indels. Notably, the majority of these mutations tend to occur outside of essential genes (**Fig. S1d**).

### Genome Signature safeguarded synthetic *E. coli* genome

The effectiveness of the Genome Signature is influenced by the number of codons present in the coded gene. Although a single gene is constrained by the number of its codons, the stability and data storage capacity of genome recording can be significantly enhanced by incorporating multiple genes (**Table S4**). For example, in the general genomes of *E. coli*, over 300 essential genes can be used as capable candidates. By dividing the genome into fragments and encoding each fragment with multiple genes—and allowing a single gene to encode multiple fragments—the same target sequence is marked by multiple genes with high QA-scores, resulting in greater stability and accuracy of genome recording. The mutated nucleotide is assigned an even higher QA-score by summing up the QA-scores of all genes that encode the same sequence (**Fig. S7**). As a result of collaboration between multiple joint genes, the accuracy of decoding and locating the mutation position is significantly improved. This collaborative effort allows for more precise identification of the mutated position in the genome (**Note S10**).

*E. coli* cells with Genome Signature encoded in ten essential genes (total of 7552 codons encoded) were constructed using standard genome editing methods (**Fig. 4a**). The Genome Signature in this *E. coli* strain consists of five unique genes, each encoding a distinct fragment, while the remaining five genes encode multiple fragments (**Table S6** and **Fig. S7**). The genome sequence is structured in five layers, with each fragment being encoded by 4-5 genes. The expression levels of the encoded genes were quantified and showed no significant differences compared to the control group (**Fig. 4b**). Furthermore, the *E. coli* cells containing the Genome Signature in multiple genes exhibited normal growth rates (**Fig. 4c**). In the computational experiments, the decoding capability was quantified by performing over a million calculations involving random mutations, including multiple substitutions, insertions, and deletions across the genome (**Fig. 4d**). Furthermore, mutations were successfully decoded from *E. coli* cells that were sequenced after undergoing treatment with Atmospheric and Room Temperature Plasma (ARTP) for random mutations generation.

**Figure 4.**
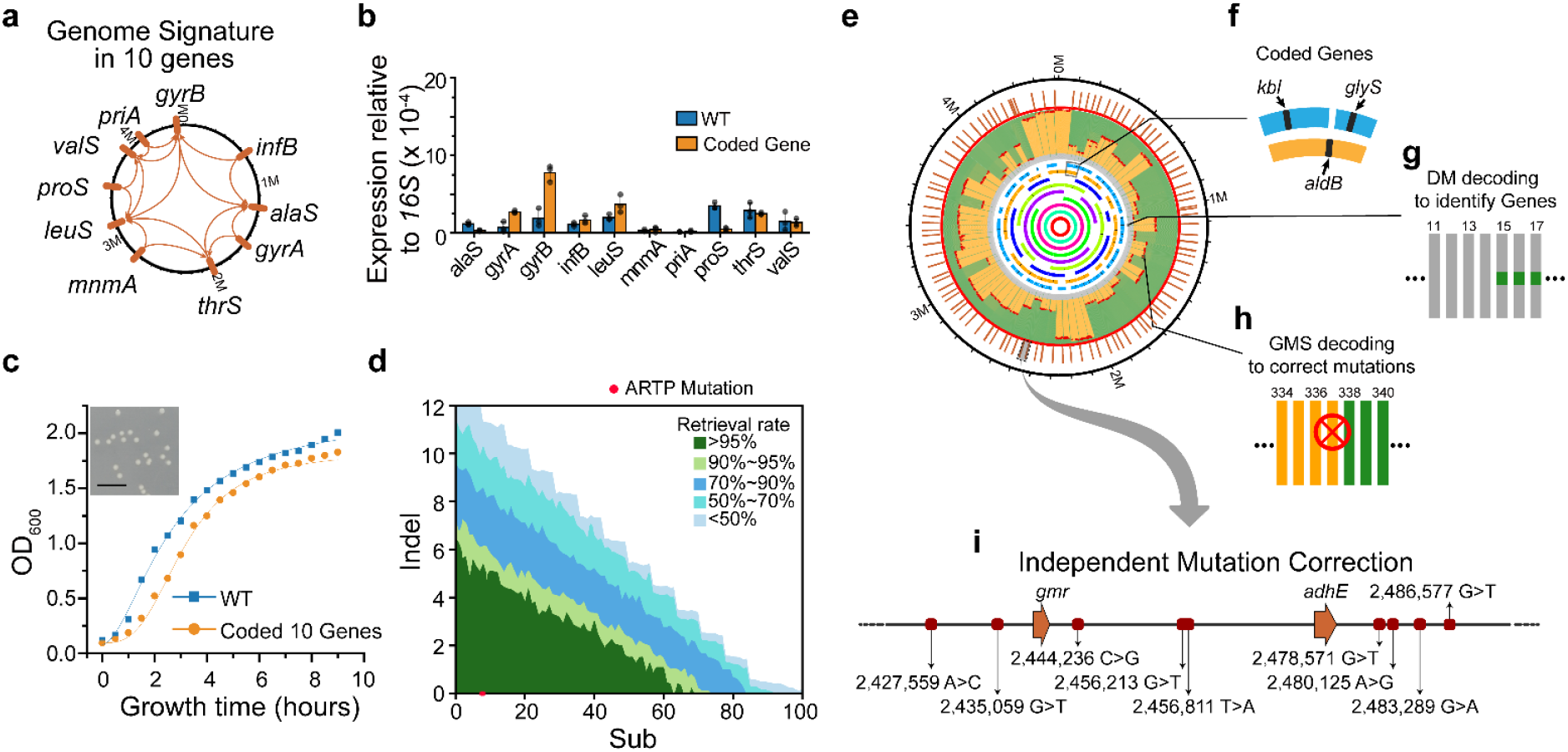
Genome Signature protected synthetic *E. coli* genome. (a) *E. coli* genome of 4,521,562 nucleotides in length with a Genome Signature that is coded in ten genes (Table S6). Each gene encodes a genome segment ranging from 622,018 to 4,521,562 nucleotides. (b) A genome with 10 coded genes is constructed by precise genome editing (Methods), and the expression levels of the ten encoded genes were quantified using quantitative PCR. (c) The cell growth is measured in standard LB medium. (d) Various combinations of random substitutions (X-axis), and indels (Y-axis) were identified in computation experiments. The success rate of correction for each mutation combination was calculated using at least 100 computations. The mutations (red dot) were successfully identified in the sequenced genome of cells that were treated with ARTP (Methods). (e) The genome of *E. coli* BL21(DE3) contains the Genome Signature encoded in a total of 100 genes, their position on the genome was marked as outer red (mutation correction) bars. A 9-layer nested structure is depicted in the center as curved color lines, representing the 10 segments that are coded within these 100 genes (Table S6). (f) Gene *kbl* codes a segment of 76,795 nucleotides (right blue curve line), *glyS* codes the downstream segment of 64,937 nucleotides (left blue curve line) and *aldB* codes segment spanning both *kbl* and *glyS* segments of 143,271 nucleotides (orange curve line). (g) All of the 100 coded genes were identified in iterative DM decoding cycles depicted as numbered gray lines. (h) A total of 700 substitution mutations were identified and corrected. A Red cross shows a successfully corrected mutation at round 337 of the GMS decoding. Orange lines indicate sequences with undetermined authenticity, while green lines represent sequences with GMS-validated authenticity. (i) Distribution of decoded Genome Signature and mutations in the local genome.

Moreover, we also tested the correction of fragment mutations that involved multiple consecutive substitutions or indels. In contrast to robust decoding of up to 20 consecutive substitutions, the successful decoding of indels was limited to only 2-3. As described above, indel causes more codon-changes than substitution (**Fig. S5a**), but indel correction can be further improved, e.g., recording more auxiliary information, such as the fragment size, in the Genome Signature. In theory, identifying fragment indels is a highly complex problem, and there are currently no practical algorithms available(*23*). However, Genome Signature has the ability to locate the positions of fragment indels on the genome (**Fig. S7**). Since fragments with severe mutations do not impact the decoding of other genome fragments, the encoding of the Genome Signature in multiple genes efficiently reduces the risk of collapse during decoding. Long-term bacterial evolution studies(*22, 24, 25*) have shown that bacterial genomes accumulate only a few mutant nucleotides each year (**Table S1**). Therefore, by encoding the Genome Signature in 10 joint genes, the genome demonstrated improved robustness.

Furthermore, a more effective Genome Signature was encoded in 100 jointly coded genes (totaling 73,557 codons), which included 11 essential and 89 nonessential genes (**Table S6**). The 100 genes formed a 9-layer nested structure (**Fig. 4e**) to achieve a high capability for mutation correction. The shortest segment of the genome coded in the gene *frwA* (834 codons) is 24,260 nucleotides, while *fecA* (775 codons) codes the entire genome consisting of 4,521,562 nucleotides. Consequently, on average, every segment of the genome sequence is coded by 7 genes. In a computational experiment (**Table S5**), the genome sequences with randomly mutated nucleotides were subjected to decoding. All 100 coded genes were identified in 100 repeated DM decoding cycles (**Fig. 4f, g**) after correcting mutations in the sequence of all coded genes in about 2 seconds. Next, hundreds of mutations outside the coded gene sequences were identified and corrected through 703 iterative GMS decoding cycles, which took approximately 136.82 hours (**Fig. 4h, i** and **Movie S1**). After the correction of 700 substitution mutations, the integrity and authenticity of the genome were successfully determined without querying any references for comparison. In another computational experiment (**Fig. S8**), a total of 97 substitutions and 10 indels were identified and corrected, resulting in the recovery of approximately 97.39% of the whole genome sequence. However, there was a failure to correct 53 substitutions and 50 indels on a single fragment of 118,126 nt. Severe mutations may overload the computation and cause a failure in decoding one fragment, but the damage can be controlled without affecting other fragments.

### Tracking the Genome Signature across Distinct Genomes

A partial sequence of a synthetic genome, particularly one designed with gain-of-function features, has the potential to be horizontally transferred into other cells(*26*), thereby raising significant biosafety concerns within the rapidly advancing field of synthetic biology. Concurrently, the unauthorized use of genomic sequences featuring unique intelligent designs poses a major issue regarding intellectual property rights in the GMO-related industry(*27*). The Genome Signature, with its high stability and distinct mechanism, can precisely trace an encoded sequence embedded within various genomic contexts. This capability allows it to determine the origin and authenticity of the sequence without the need for sequence comparison against references, which is particularly valuable for confirming its ownership in legal disputes.

To demonstrate the feasibility, a 7,035 nt fragment of the *E. coli* Nissle 1917 genome was encoded with the Genome Signature in the gene *iutA* (**Fig. 5a**). *E. coli* Nissle 1917 is a probiotic strain that is widely used in biomedicines and food production(*28*). The Nissle cell was constructed using precise genome editing techniques and grew normally (**Fig. 5b**). To mimic the horizontal transfer of genomic material, this Nissle genomic fragment containing the Genome Signature was inserted into three different locations within the *E. coli* BL21 genome. The genetically invaded BL21 cells were cultured, and total genomic DNA was collected. Subsequently, the collected DNA samples were subjected to high-accuracy long-read sequencing using the PacBio HiFi platform, which has a sequencing error rate lower than 0.1% (**Methods**). A total of 323,846 PacBio long reads, each with a length of over 2 Kbp, were collected and subjected to individual Genome Signature decoding. All 312,631 long reads were processed, and nucleotide mutation correction and authenticity determination were carried out using approximately 7.3E+14 hash calculations. Ultimately, 40 sequences carrying the full 7,035-nt invaded sequence were successfully determined (**Fig. 5c**), while only a partial invaded sequence (coded *iutA*) could be determined in 144 sequences. To assess the limitations of Genome Signature, an Australian lungfish genome, which is regarded as one of the largest sequenced genomes to date(*29*), was utilized to track migrated genomic sequences. Initially, a single, comprehensive genome sequence was constructed by concatenating all 14 chromosomes (GCA_016271365.1) of the lungfish, resulting in a final size of 34,557,647,948 nt (**Fig. 5d**). The Nissle fragment was inserted at base position 18,144,212,270 within this composite genome. Remarkably, even in the context of such high sequence complexity, the full invaded fragment was successfully determined through approximately 11.7 billion Hash calculations.

**Figure 5.**
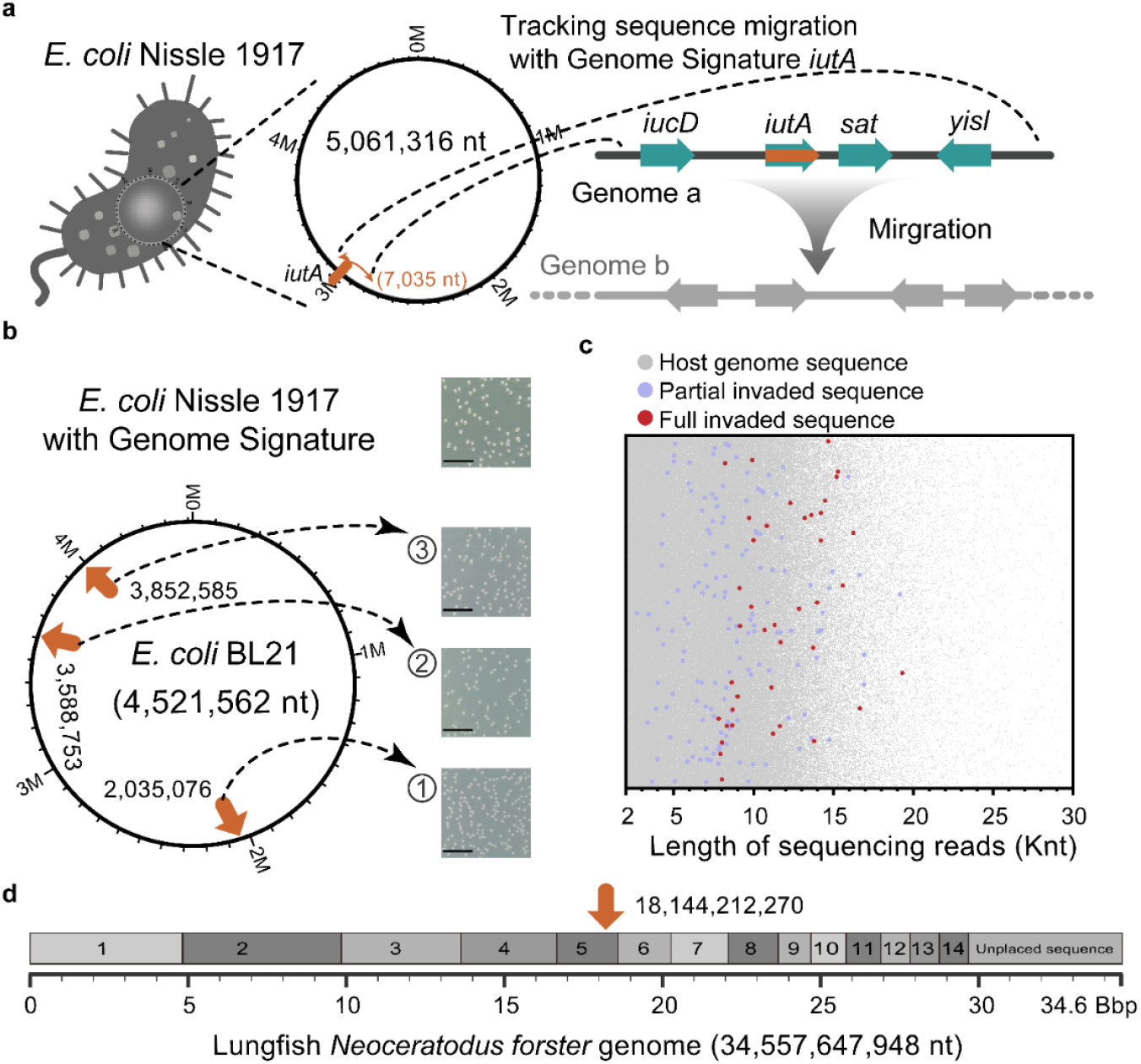
Tracking Genome Signature across Distinct Genomes. (a) One fragment of 7035 nt of *E. coli* Nissle 1917 with Genome Signature in one gene of *iutA*. (b) Nissle *iutA* fragment was inserted in the genome of *E. coli* BL21 at different locations (1: 2,035,076; 2: 3,588,753; 3: 3,852,585), *E. coli* cells cultured on standard LB solid medium. (c) These *E. coli* BL21 strains were sequenced using the PacBio platform, and long reads exceeding 2 Kbp in length were directly decoded. Reads with a determined full *Nissle* fragment are indicated by red dots, while reads with only the determined coded *iutA* gene but not the full length of the fragment are represented by purple dots. The y-axis indicates the position of the sequencing read on the genome. (d) Nissle coded *iutA* fragment (inserting at 18,144,212,270) was successfully determined in a giant genome sequence constructed by jointing all 14 lungfish chromosomes (GCA_016271365.1), a total of 34,557,647,948 nt in length.

Unlike traditional DNA tag or watermark frameworks, Genome Signatures possess the capability to both identify the presence of their own and directly assess the integrity and authenticity of the entire genomic sequence. Consequently, Genome Signatures could serve as a powerful tool for monitoring the contamination of synthetic sequences in natural organisms and detecting the unauthorized use of intelligently designed genomic materials.

## Discussion

Today, genomic sequence information is indeed considered a form of Big Data, with global databases storing dozens of exabytes(*30*). Furthermore, the field is anticipated to experience even more rapid growth in the future, presenting additional hurdles for existing genome characterization methods. The high-integrity storage of genomic data in centralized public databases, essential for genomic analysis, has been hindered by various technical, ethical, and legal concerns. This powerful Genome Signature platform has the potential to revolutionize the centralized storage of crucial genetic information. Instead of relying on a centralized database, the platform stores the information for integrity, identification, specification, and intellectual property rights protection directly in the genome itself. With the multitude of challenges our society faces due to the rapid development of GMOs, Genome Signature promises viable solutions and can play a significant role in advancing synthetic biology for both scientific research and real-world applications.

Genome Signature can be encrypted into a genome in living cells while also preserving the integrity of the genome structure and causing no changes to the size of the genome sequence. The inherent connection between coded genes and the genome sequence diminishes the likelihood of damage or loss in genome-wide Genome Signatures. In this study, we developed living bacteria *E. coli* cells with a robust Genome Signature in as many as ten genes, but the method can easily be scaled up to include more genes. Genome Signatures represent a novel platform for directly and accurately determining genome integrity and verifying the authenticity of genomic sequences, all without the need for reference-based sequence comparisons. Unlike conventional DNA tag frameworks, Genome Signatures have the potential to revolutionize the administration of synthetic genomic materials by offering a reference-free approach. Just like the stability and security achieved in blockchain technology, once encrypted, the algorithmic connection and digital data encrypted inside essential genes cannot be tampered with. Furthermore, the combination of directed hashing in the Genome-CHM algorithm gives Genome Signatures the ability to provide certification. The use of authentication keys is able to ensure high security. As a result, Genome Signatures have the potential to achieve a high level of biosafety and can be a powerful tool for protecting intellectual property and safeguarding GMOs. The exponential expansion of synthetic genomes has spurred widespread concerns regarding their potential impact on multiple facets of society, with a special emphasis on environmental safety. Genome Signatures can address these concerns by enabling independent monitoring, without the need for reference-based sequence comparisons, of potential contamination of synthetic genome sequences in natural systems.

## Methods

### Genome editing

To insert Genome Signature into the genome of *E. coli*, we prepared 11 linear recombination engineering templates to replace the corresponding wild-type gene sequence of Genome Signature. The DNA sequence of the linear recombination template includes the upstream 500 bp sequence adjacent to the Genome Signature in the genome (the upstream homology arm), the Genome Signature sequence, the ampicillin-resistant (AmpR) gene sequence used as a screening marker, and the downstream 500 bp sequence adjacent to Genome Signature in the genome (the downstream homology arm). Genome Signature sequences and the AmpR sequence used as a screening marker were synthesized by Liuhe BGI Tech Co. Ltd (Beijing, China). The upstream and downstream homology arms were amplified from the genome in a 50 μl PCR reaction using PrimeSTAR® Max DNA Polymerase (Takara, China), respectively. Overlapping PCR was used to prepare the linear recombination template for gene editing. These fragments were then used to transform competent *E. coli* BL21(DE3) S-ABP pTKRED(*31*), which is collectively referred to as *E. coli* BL21(DE3) in this article, according to the method proposed by Kuhlman *et*.*al(32)*, see **Table S7** for the genome of *E. coli* BL21(DE3).

We used the genome editing methods based on pEcCas/pEcgRNA(*33*) for simulating the migration of a fragment of 7,035 nt watermarked by Genome Signature *iutA* from *E. coli* Nissle 1917 into *E. coli* BL21. gRNA target for each of these positions was designed on https://chopchop.cbu.uib.no/ to construct of pECgRNA-target plasmid. The linear recombination template for gene editing includes the 500 bp upstream homology arm, the migration fragment, and the 500 bp downstream homology arm. The sequence of the homology arm was amplified from the genome of *E. coli* BL21 in a 50 μl PCR reaction using PrimeSTAR® Max DNA Polymerase, respectively. The migration fragment was amplified from the genome of *E. coli* Nissle 1917 in a 50 μl PCR reaction too. The homology arm DNA fragment and migration fragment were assembled by overlapping PCR to prepare the linear recombination template for gene editing. Competent cells for 100 μl were mixed with 1000 ng of linear recombination template and 100 ng pEcgRNA-target plasmids. The resulting mixture was then electroporated. Cells were resuspended in 1 ml SOB medium after electroporation. The cells were allowed to recover at 37°C for 1 h and then plated on LB agar plates with 50 μg/ml Kanomycin, and 100 μg/ml spectinomycin incubated overnight at 37°C.

### Cell culture

The growth of *E. coli* BL21(DE3) cell, cell coded in Genome Signature *valS*, and cell coded in ten joint Genome Signatures was tested in LB medium (10 g/l tryptone, 5 g/l yeast extract, 10 g/l NaCl). First, the cells were cultured in LB medium overnight at 37°C with shaking at 220 r.p.m. The overnight cultured cells were added to LB medium in the ratio of 1:100, respectively, and cultured at 37°C with shaking at 220 r.p.m. The optical density (OD) was measured at 565 nm every 0.5 h using a DEN-1B Densitometer (Keison products, Germany).

### Quantitative PCR

Quantitative PCR (qPCR) was used to measure the expression of ten genes (*infB, alaS, gyrA, thrS, mnmA, leuS, proS, valS, gyrB*, and *priA*) in *E. coli* BL21(DE3) cell and cell coded in ten joint Genome Signatures. The cells were cultured in LB medium overnight at 37°C with shaking at 220 r.p.m. Total RNA was extracted from growth-saturated cells using TRIzol^®^ reagent (Invitrogen, USA) according to the manufacturer’s protocol. HiScript II Q RT SuperMix for qPCR (+gDNA wiper) (Vazyme, China) was used to remove genomic DNA contamination in the sample and reverse transcript the sample. qPCR reaction was prepared with AceQ qPCR SYBR Green Master Mix (Vazyme, China) and performed in 20 μl reaction mixtures using a QuantStudio6 Flex Real-Time PCR System (Thermo Fisher Scientific, USA) following the manufacturer’s protocol. We selected *16S* RNA as a reference gene. Primers were listed in **Table S8**. Δ*C_T_* = *C_T_^gene^*-*C_T_^16S^*, so gene expression relative to *16S* RNA is 2^−Δ*CT*^.

### Mutagenesis using Atmospheric and Room Temperature Plasma (ARTP)

Fresh single colony of *E. coli* BL21(DE3) pTKRED was cultured in 3 ml of LB medium with 100 μg/ml spectinomycin overnight at 30°C. The next day, the overnight cells were transferred to LB medium with 100 μg/ml spectinomycin at 1:100. Once the OD (*λ*= 565 nm) of the culture reached 0.6, cells were diluted 10 times by LB medium, 20 μl diluted culture was treated by atmospheric and room temperature plasma (ARTP) using an ARTP mutation system (ARTP-IIS, Tmaxtree Biotechnology Co, Ltd, Wuxi, China) with the following parameters: (1) the radio frequency power input was 120 W; (2) the flow of pure helium was 10 standard liters per min; (3) the distance between the plasma torch nozzle exit and the slide was 2 mm; and (4) the treatment times were 120 s. After treatment, the slide was washed with 1 ml of LB medium in a 1.5 mL falcon tube, and cultivated at 30°C and 220 r.p.m. for 1 h. Then the culture medium was gradient diluted and coated onto an LB plate. A single colony was picked to 3 ml of LB medium with 100 μg/ml spectinomycin and culture at 30°C overnight. The next day, the cells were transferred to LB medium with 100 μg/ml spectinomycin at 1:100. Cells were collected by centrifugation once the OD (*λ* = 565 nm) of the culture reached 0.6 and washed twice with 1 ml of PBS buffer (PH7.4 slightly), then the bacteria were frozen by liquid nitrogen and send for sequencing.

### Genome long-read sequencing

Single colonies of *E. coli* BL21 with migrated fragments at different locations were selected and cultured in 3mL LB medium overnight respectively. The next day, the cells were mixed and transferred to 3L LB medium at 1:1000. Cells were collected by centrifugation once the OD (λ = 565 nm) of the culture reached 0.6 and washed twice by 50 mL of PBS buffer (PH7.4 slightly), then the bacteria was frozen by liquid nitrogen and sent for PacBio sequencing. Each sequencing reads were decoded to find the migrated fragment.

### The giant genome of lungfish

The genome data of the lungfish *Neoceratodus forsteri* (Australian lungfish) is from Genbank (GCA_016271365.1). Its genome size is 34,557,647,948 nt after removing 129 N. The final linear genome is obtained by connecting 15 fragments one after the other according to the following sequence: chromosome 1-14 and unplaced sequence fragment. Linear genome carrying a migration fragment containing the Genome Signature *iutA* serves as the decoding input.

### Computing and Statistics

All encoding and decoding data were obtained from the following machines: 32Gbyte of memory, 64 cores, 2.50 GHz Intel Xeon CPU E5-2682 v4; 16Gbyte of memory, 8 cores, Intel Xeon Processor (Skylake, IBRS); 48Gbyte of memory, 12 cores, 2.67 GHz Intel Xeon CPU x5640. The total amount of Hash calculations for this work exceeds 1.30E+16. A list of computing details and statistical tests can be found in **Table S5**.

## Acknowledgments

This work was supported by grants from the National Key R&D Program of China (Grant No. 2020YFA0712104). We would like to express our gratitude to Professor Yan Zhang for insightful discussions. Additionally, thanks to Xiaoyan Wu and Mingyi Qi for their unwavering support throughout the project.

## Author contributions

Z.G.W and J.S.C contributed equally to this work. Z.G.W. and Q.L. conducted biochemical experiments and data analysis, prepared the figures, and contributed to the manuscript preparation. Z.G.W. and J.J.L. contributed to the genome editing experiments; J.S.C., H.X., J.W.L., C.G., C.Y.J., and Y.Y.L. wrote the code and conducted all the computation experiments. Y.G.K. contributed to the population genetics study design, figure preparation, and interpretation of the results; H.Q., J.S.C., Y.G.K., and Z.G.W. prepared the manuscript together. H.Q. and J.S.C. supervised the whole project, led the development of the algorithm and designed all experiments.

## References

1. W. Weber, M. Fussenegger, Emerging biomedical applications of synthetic biology. Nat. Rev. Genet. 13, 21–35 (2011).

2. G. H. Moe-Behrens, R. Davis, K. A. Haynes, Preparing synthetic biology for the world. Front. Microbiol. 4, 5 (2013).

3. E. L. Rylott, N. C. Bruce, How synthetic biology can help bioremediation. Curr Opin Chem Biol 58, 86–95 (2020).

4. J. Ke, B. Wang, Y. Yoshikuni, Microbiome Engineering: Synthetic Biology of Plant-Associated Microbiomes in Sustainable Agriculture. Trends Biotechnol. 39, 244–261 (2021).

5. X. Tan, J. H. Letendre, J. J. Collins, W. W. Wong, Synthetic biology in the clinic: engineering vaccines, diagnostics, and therapeutics. Cell 184, 881–898 (2021).

6. D. Chakravarti, W. W. Wong, Synthetic biology in cell-based cancer immunotherapy. Trends Biotechnol. 33, 449–461 (2015).

7. A. Aguilera, T. Garcia-Muse, Causes of genome instability. Annu. Rev. Genet. 47, 1–32 (2013).

8. J. W. Lee, C. T. Y. Chan, S. Slomovic, J. J. Collins, Next-generation biocontainment systems for engineered organisms. Nat. Chem. Biol. 14, 530–537 (2018).

9. T. M. Ghaly et al., Discovery of integrons in Archaea: Platforms for cross-domain gene transfer. Sci Adv 8, eabq6376 (2022).

10. F. Gould et al., Toward product-based regulation of crops. Science 377, 1051–1053 (2022).

11. D. G. Gibson et al., Creation of a bacterial cell controlled by a chemically synthesized genome. Science 329, 52–56 (2010).

12. K. Wang, D. de la Torre, W. E. Robertson, J. W. Chin, Programmed chromosome fission and fusion enable precise large-scale genome rearrangement and assembly. Science 365, 922–926 (2019).

13. J. Luo, X. Sun, B. P. Cormack, J. D. Boeke, Karyotype engineering by chromosome fusion leads to reproductive isolation in yeast. Nature 560, 392–396 (2018).

14. J. E. Gallegos, D. M. Kar, I. Ray, I. Ray, J. Peccoud, Securing the exchange of synthetic genetic constructs using digital signatures. ACS Synth. Biol. 9, 2656–2664 (2020).

15. I. Hafeez, A. Khan, A. Qadir, DNA-LCEB: a high-capacity and mutation-resistant DNA data-hiding approach by employing encryption, error correcting codes, and hybrid twofold and fourfold codon-based strategy for synonymous substitution in amino acids. Med Biol Eng Comput 52, 945–961 (2014).

16. Y. Erlich, D. Zielinski, DNA Fountain enables a robust and efficient storage architecture. Science 355, 950–954 (2017).

17. H. H. Wang et al., Programming cells by multiplex genome engineering and accelerated evolution. Nature 460, 894–898 (2009).

18. G. Kudla, A. W. Murray, D. Tollervey, J. B. Plotkin, Coding-sequence determinants of gene expression in Escherichia coli. Science 324, 255–258 (2009).

19. M. Liss et al., Embedding permanent watermarks in synthetic genes. PLoS ONE 7, e42465 (2012).

20. Q. Liu, P. Wang, J. Cui, H. Qi, paper presented at the 2020 Eighth International Conference on Advanced Cloud and Big Data (CBD), 2020.

21. N. Memarsadeghi, NASA computational case study: Golomb rulers and their applications. Compos Sci Technol 18, 58–62 (2016).

22. D. Leon, S. D’Alton, E. M. Quandt, J. E. Barrick, Innovation in an E. coli evolution experiment is contingent on maintaining adaptive potential until competition subsides. PLoS Genet. 14, e1007348 (2018).

23. W. J. P. Spee, J. H. Weber, Bounds on the Maximum Cardinality of Indel and Substitution Correcting Codes. IEEE Transactions on Molecular, Biological, and Multi-Scale Communications 10, 349–358 (2024).

24. H. Lee, E. Popodi, H. Tang, P. L. Foster, Rate and molecular spectrum of spontaneous mutations in the bacterium Escherichia coli as determined by whole-genome sequencing. Proc Natl Acad Sci USA 109, E2774–2783 (2012).

25. P. R. Reeves et al., Rates of mutation and host transmission for an Escherichia coli clone over 3 years. PLoS One 6, e26907 (2011).

26. P. Edelaar, D. I. Bolnick, Non-random gene flow: an underappreciated force in evolution and ecology. Trends Ecol. Evol. 27, 659–665 (2012).

27. S. Mueller, F. Jafari, D. Roth, A covert authentication and security solution for GMOs. BMC Bioinform 17, 389 (2016).

28. L. Grozdanov et al., Analysis of the genome structure of the nonpathogenic probiotic Escherichia coli strain Nissle 1917. J Bacteriol 186, 5432–5441 (2004).

29. A. Meyer et al., Giant lungfish genome elucidates the conquest of land by vertebrates. Nature 590, 284–289 (2021).

30. Z. D. Stephens et al., Big Data: Astronomical or Genomical? PLoS Biol. 13, e1002195 (2015).

31. Y. Wu et al., Efficient In Vitro Full-Sense-Codons Protein Synthesis. Advanced Biology, 2200023 (2022).

32. T. E. Kuhlman, E. C. Cox, Site-specific chromosomal integration of large synthetic constructs. Nucleic Acids Res. 38, e92 (2010).

33. Q. Li et al., A modified pCas/pTargetF system for CRISPR-Cas9-assisted genome editing in Escherichia coli. Acta Biochim. Biophys. Sin. 53, 620–627 (2021).

